# Elucidating the molecular determinants of Aβ aggregation with deep mutational scanning

**DOI:** 10.1101/662213

**Authors:** Vanessa E. Gray, Katherine Sitko, Floriane Z. Ngako Kameni, Miriam Williamson, Jason J. Stephany, Nicholas Hasle, Douglas M. Fowler

**Affiliations:** Department of Genome Sciences, University of Washington, Seattle, WA, USA; Department of Bioengineering, University of Washington, Seattle, WA, USA; Genetic Networks Program, CIFAR, Toronto, ON, Canada

## Abstract

Despite the importance of Aβ aggregation in Alzheimer’s disease etiology, our understanding of the sequence determinants of aggregation is sparse and largely derived from *in vitro* studies. For example, *in vitro* proline and alanine scanning mutagenesis of Aβ_40_ proposed core regions important for aggregation. However, we lack even this limited mutagenesis data for the more disease-relevant Aβ_42_. Thus, to better understand the molecular determinants of Aβ_42_ aggregation in a cell-based system, we combined a yeast DHFR aggregation assay with deep mutational scanning. We measured the effect of 791 of the 798 possible single amino acid substitutions on the aggregation propensity of Aβ_42_. We found that ~75% of substitutions, largely to hydrophobic residues, maintained or increased aggregation. We identified 11 positions at which substitutions, particularly to hydrophilic and charged amino acids, disrupted Aβ aggregation. These critical positions were similar but not identical to critical positions identified in previous Aβ mutagenesis studies. Finally, we analyzed our large-scale mutagenesis data in the context of different Aβ aggregate structural models, finding that the mutagenesis data agreed best with models derived from fibrils seeded using brain-derived Aβ aggregates.

## Introduction

Protein aggregation affects all known organisms from bacteria to humans and is implicated in a number of human diseases. Decades of genetic, biochemical and epidemiological work suggests that aggregation of the amyloid β (Aβ) peptide is related to the incurable neurodegeneration associated with Alzheimer’s disease (Hardy and Selkoe 2002; Lesné *et al.* 2008; Bertram and Tanzi 2008; Shankar *et al.* 2009; Masters and Selkoe 2012; Hardy 2017). Aβ peptide is generated by post-translational cleavage of the transmembrane amyloid β precursor protein at variable positions to produce peptides that range from 38 to 43 amino acids in length. The most aggregation-prone form of Aβ is 42 amino acids long (Aβ_42_), though Aβ_40_ is present at higher concentrations in human cerebrospinal fluid (Jarrett *et al.* 1993; Iwatsubo *et al.* 1994; Dahlgren *et al.* 2002). The aggregation of Aβ begins with a shift in equilibrium from soluble monomers to oligomers, and these oligomers may nucleate amyloidogenesis (Matsumura *et al.* 2011; Barz *et al.* 2018). In Alzheimer’s disease, Aβ fibrils accumulate in the extracellular space forming the major component of amyloid plaques, a defining feature of the disease.

Despite the importance of Aβ aggregation in Alzheimer’s disease etiology, our understanding of the sequence determinants of aggregation is sparse and largely derived from *in vitro* studies. In the past decade, several assays based on the budding yeast *S. cerevisiae* have been used to study protein aggregation (Bagriantsev and Liebman 2006; Haar *et al.* 2007; Caine *et al.* 2007; Morell *et al.* 2011; D’Angelo *et al.* 2013). Notably, a growth-based assay that separates toxicity from aggregation offers a way to investigate how changes in Aβ sequence impact aggregation propensity (Morell *et al.* 2011) (**Figure 1A**). In this assay, Aβ is cytoplasmically localized to eliminate its aggregation-associated toxicity (Treusch *et al.* 2011; D’Angelo *et al.* 2013). To link Aβ aggregation to yeast growth, Aβ is fused to an essential protein, dihydrofolate reductase (DHFR) via a short peptide linker. The result is that DHFR activity depends on the solubility of Aβ. Thus, upon treatment with the competitive DHFR inhibitor methotrexate, yeast expressing soluble Aβ variants grow rapidly, whereas yeast expressing aggregating Aβ variants grow slowly.

**Figure 1A-D.**
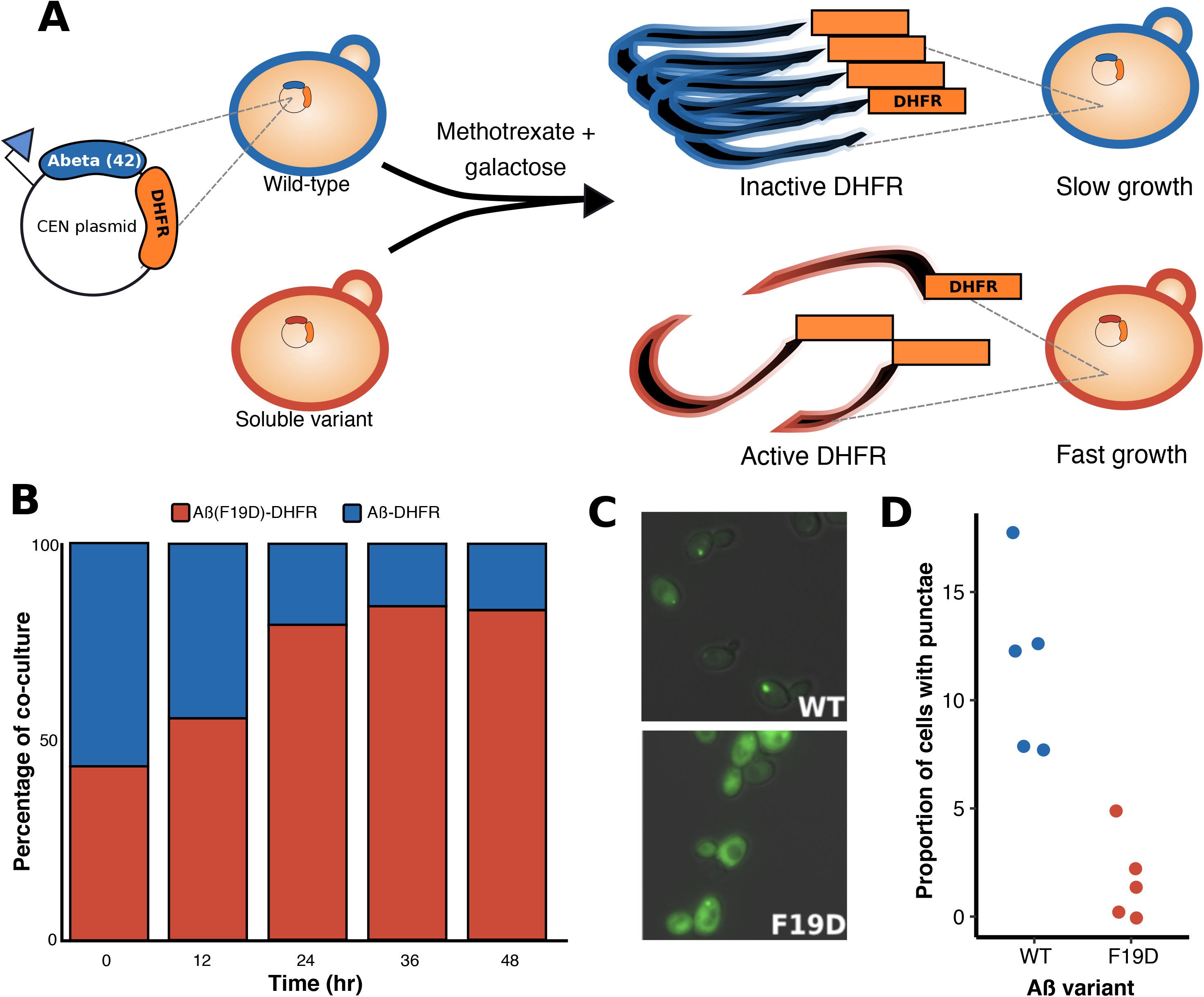
A yeast-based aggregation assay distinguishes between soluble and aggregation-prone variants of Aβ. A schematic of the assay shows plasmid-based expression of Aβ-DHFR and a nonaggregating variant of Aβ fused to DHFR lead to slow and fast yeast growth in the presence of methotrexate, respectively (**A**). A stacked bar graph shows the percentage of Aβ-DHFR and Aβ(F19D)-DHFR (less aggregation prone variant) in co-culture (y-axis) every 12 hours for 48 hours (x-axis; **B**). Fluorescence light microscopy shows the aggregation patterns of Aβ-GFP (WT) and Aβ(F19D)-GFP (F19D) 16h after induction of expression (**C**). A bar graph shows the proportion of yeast cells with punctae (y-axis) in five fluorescence microscopy images of Aβ-DHFR (WT) or Aβ(F19D)-DHFR (F19D; x-axis; **D**).

Mutagenesis can elucidate the role of individual residues in protein aggregation. For example, *in vitro* proline (Williams *et al.* 2004) and alanine (Williams, Shivaprasad, and Wetzel 2006a) scanning mutagenesis of Aβ_40_ revealed core regions important for aggregation. However, we lack even this limited mutagenesis data for the more disease-relevant Aβ_42_ and, so far, the majority of mutagenesis studies have been performed *in vitro*.

Thus, to fully understand the molecular determinants of Aβ_42_ aggregation in a cell-based system, we combined the yeast DHFR aggregation assay with deep mutational scanning (Araya and Fowler 2011; Fowler and Fields 2014; Fowler *et al.* 2014) to measure the effect of 791 of the possible 798 single amino acid substitution on the aggregation propensity of Aβ_42_. We used high-throughput DNA sequencing to track the frequency of each Aβ_42_ variant during the selection, enabling us to assign a solubility score to every variant. We present the first large-scale, cell-based mutational analysis of Aβ, illuminating the physicochemical properties of amino acids that abrogate, promote or do not effect Aβ aggregation. Of 791 single amino acid Aβ variants we evaluated, ~75% maintained or increased aggregation. In addition, we identified 11 positions at which substitutions, particularly to hydrophilic and charged amino acids, disrupted Aβ aggregation. These critical positions were similar but not identical to critical positions identified in previous Aβ mutagenesis studies. Finally, we analyzed our large-scale mutagenesis data in the context of different Aβ aggregate structural models, finding that some structures were plausible whereas others were not.

## METHODS

### Library construction

The library was cloned using *in vivo assembly* (García-Nafría *et al.* 2016). First, a forward primer containing a 5’ homology region, an NNK codon, and a 3’ extension region was designed for each codon in Aβ_42_. The homology and extension regions were at least 15 nucleotides in length and had melting temperatures greater than 55C. Reverse primers were the reverse complement of the 5’ homology region.

A separate PCR reaction was performed for each codon. These reactions contained 40 ng template (p416GAL1-Aβ-DHFR) and 10 μM forward and reverse primers (IDT, custom oligos) in a total reaction volume of 30 μL. The following cycling conditions were used: 95C 3min, 8× [98C 20 sec, 60C 15 sec, 72C 9 min], 72C 9 min. After PCR, 7.5 μL of each product was run on a 1.5% agarose gel for 30 min at 100V to check for a single product. The remaining 22.5 μL aliquots of product were each digested for an hour at 37C with 0.6 μL of DpnI (NEB, R0176S). After digestion, 4 μL of each linear product was transformed into a 50 μL of TOP10F Chemically Competent *E. coli* (ThermoFisher, C303003) according to manufacturer’s instructions, with the following modifications: the protocol was done in a 96 well plate, and cells were recovered in a total volume of 200 μL SOC. After recovery, cells were transferred to a deep well plate with 1.6-1.8 mL of ampicillin LB and shaken overnight. To estimate colony count, 50 μL of culture was plated on an LB + ampicillin agar plate. Deep well plates and agar plates were incubated at 37C overnight. After incubation, all 42 deep well plate cultures were combined and subject to a midiprep (Sigma, NA0200).

### 4.3.2 Plasmids, yeast strains and growth conditions

To create a galactose-regulated Aβ-DHFR expression system, we cloned the human Aβ_42_ coding sequence into the SpeI and HindIII sites of p416 (URA3, GAL1 promoter, CEN) (Mumberg, Mu◻ller, & Funk, 1994). Aβ-GFP variants were cloned using the same scheme. All Aβ variants were cloned into p416 and transformed in W303 strain (MATa/MATα {leu2-3,112 trp1-1 can1- 100 ura3-1 ade2-1 his3-11,15} [phi+]). Cells were grown at 30C in synthetic complete (SC) media lacking uracil and supplemented with 2% glucose.

### Methotrexate selection assay

Transformed yeast were inoculated into 5 mL (low-throughput) or 300 mL (deep mutational scan) of C-Ura, 2% glucose media, grown in a rotating/shaking, 30C incubator overnight and then transferred to 5 mL or 300 mL 2% raffinose media to remove the glucose repression acting on the *gal1* promoter. After two hours in 2% raffinose, yeast were back-diluted to an OD of 0.01 into 5 mL or 300 mL 2% galactose to induce Aβ_42_-DHFR expression in the presence or absence of 80 μΜ methotrexate (TCI America, M-1664), 1 mM sulfanilamide (Sigma, S-9251). For 300 mL experiments, wherein yeast with aggregating and nonaggregating variants were grown in co-culture, the two strains were inoculated at equally densities. In 5 mL experiments, yeast growth was measured using a spectrophotometer that detects 660 nm wavelengths over a 48h course. In competition experiments, 10 OD units of yeast were collected from 300 mL cultures every 12h, spun down, concentrated and stored in −80C. At the end of the experiment, frozen yeast were thawed and then their plasmids were extracted using a DNA Clean and Concentrator kit (Zymo Research, D4013). Extracted plasmids were prepped and sequenced using Sanger sequencing. The following equation was used to calculate doubling times from two time points: (Log_10_(OD_T2_/OD_T1_)/ Log_10_(2))/ ΔT, where OD represents the optical density at 600nm at a time point (T).

### Library preparation for high-throughput sequencing

Plasmids were extracted from yeast with Yeast Plasmid Miniprep 1 kit (Zymo Research, D-2001). Library fragments were amplified in 17 PCR cycles using primers specific to DNA sequences that flank Aβ-DHFR in p416, and sequenced by an Illumina NextSeq sequencer using paired-end reads (**Table S1**).

### Variant effect analysis

Enrich2 was used to calculate solubility scores for each Aβ variant from sequencing fastq files (Rubin *et al.* 2017). The Enrich2 pipeline calculates a variant’s score in three steps. First, a variant’s normalized frequency ratios are tabulated for each timepoint by dividing the frequency of a variant’s sequencing reads by all mapped reads and normalizing by the wild-type frequency ratio. Sequencing reads were stringently filtered for quality by requiring each base have a Phred score greater than 20 and no uncalled bases. Second, a weighted linear least squares regression line is fit to the normalized frequency ratios across time. Third, the slope of the regression line is calculated and averaged across the three replicates. This averaged slope reflects a variant’s aggregation propensity. Solubility scores below 0 denote variants that are more aggregation-prone than wild-type, whereas scores above 0 indicate that a variant has increased solubility compared to wild-type.

### Classifying Aβ variants using synonymous mutations

Variant classifications (*i.e.,* WT-like, more aggregation-prone, more soluble) were assigned using the distribution of 39 synonymous mutations from the deep mutational scan. We define WT-like as any variant with a score within ±2 standard deviations of the synonymous variant mean [−0.26,0.39]. A variant is more-aggregation prone than wildtype if its score is greater than 0.39 or more soluble if its score is lower than −0.26.

### Data and code availability

<u>Raw sequencing data will be made available upon publication in the NCBI GEO database. Code is available at</u> https://github.com/FowlerLab/amyloidBeta2019.

## Results

First, we verified that the DHFR-based yeast aggregation assay could differentiate between aggregating wild type Aβ (Aβ_WT_) and a nonaggregating (Aβ_F19D_) variant (Morell *et al.* 2011). As expected, in a mixed culture treated with methotrexate, Aβ_F19D_ outcompeted Aβ_WT_ (**Figure 1B**). We used fluorescence microscopy of Aβ-GFP fusions to confirm that 49% of yeast expressing Aβ_WT_-GFP had cytoplasmic punctae compared to 6% of cells expressing Aβ_F19D_-GFP (**Figure 1C-D**). Thus, we concluded the assay could be used in a deep mutational scan to measure the aggregation propensity of variants of Aβ.

Using this assay, we conducted a deep mutational scan of Aβ that yielded solubility scores for 791 single amino acid variants, representing 99.1% of the possible single variants. Solubility scores were calculated by taking the weighted least squares slope of each variant’s frequency ratios across six time points. (**see Methods**). The slopes from each replicate were well correlated (Pearson’s R 0.78 to 0.92; **Figure 2A, Figure S1A**). Replicate slopes were averaged and normalized to produce final solubility scores such that wild-type had a solubility score of zero (**Table S2**). Positive solubility scores indicated less aggregation and negative scores indicated increased aggregation.

**Figure 2A-F.**
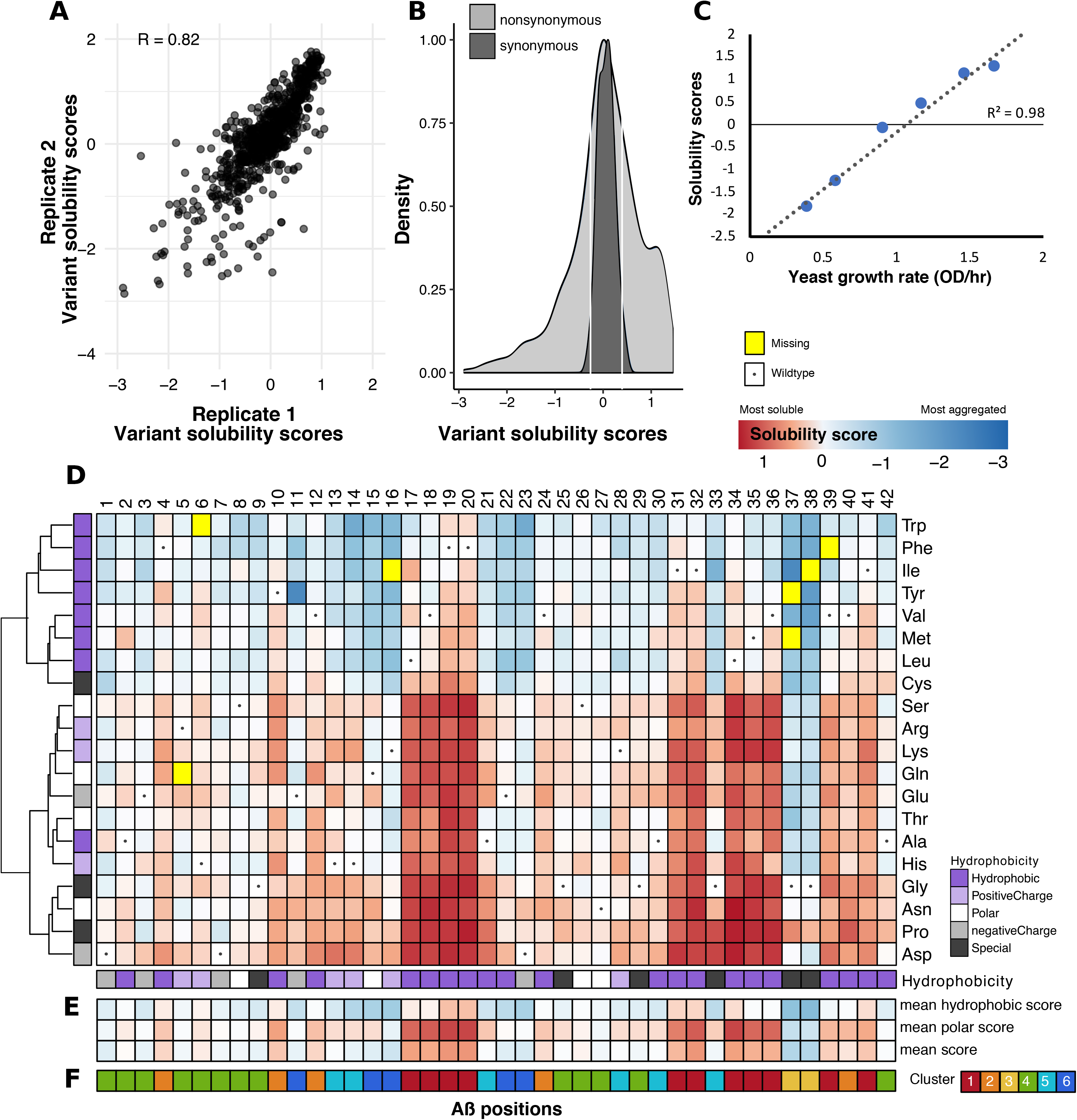
Solubility scores for 791 Aβ variants. Solubility scores reliably measure the effects of Aβ sequence on aggregation propensity. A scatter plot shows the correlation between two of three biological replicates that were averaged to yield final solubility scores (**A; Figure S1A**). The distribution of solubility scores (x-axis) of synonymous variants was used to determine cutoffs that define variants that are wild-type-like or more/less aggregation-prone than wild-type. The density plot shows distributions of nonsynonymous (light gray) and synonymous (dark gray) variants and the white lines show the lower (−0.26) and upper (0.39) bounds for wild-type-like mutations (**B**). The scatterplot shows the correlation between our solubility scores (y-axis) and a low-throughput yeast growth assay that measured yeast growth rate as a proxy for Aβ solubility (**C; Figure S1B**). The heatmap shows the effect of 791 Aβ variants on solubility with Aβ positions on the x-axis and mutant amino acids on the y-axis. A variant’s color denotes its solubility: red is most soluble, white is wild-type-like and, dark blue is most aggregated, whereas yellow variants are missing from our variant library and dots denote the wild-type amino acid at a given position. The annotation tracks on the x- and y-axes display the hydrophobicity of each wild-type and mutant amino acid, respectively. The heatmap’s y-axis has been re-ordered using hierarchical clustering on the solubility score vectors (**D**). For each position, the mean solubility score at each position is depicted using the same color scheme as the main heatmap. Additionally, the mean solubility scores for all hydrophobic and polar substitutions are shown (**E; Figure S2A**). Heirarchical clustering on the x-axis yielded 6 distinct clusters: 1 (red), 2 (orange), 3 (yellow), 4 (green), 5 (light blue), and 6 (dark blue; **F**; **Figure S2B-C**).

Solubility scores ranged from −2.38 (most aggregating) to 1.45 (most soluble). The mean (median) solubility score for all variants was 0.09 (0.08), which was similar to the solubility scores of the 39 synonymous variants in our library (mean: 0.06; median: 0.08). Because we did not expect synonymous variants to affect aggregation propensity, we used their distribution of scores to identify WT-like variants (**Figure 2B**). In total, we found that 344 (43.4%) of Aβ variant scores were within two standard deviations of the synonymous score mean and thus had WT-like effects (WT-like range: [-0.26,0.39]). Additionally, we found 246 (31.1%) variants to be more aggregation-prone than Aβ_WT_ and 201 (25.4%) variants to be more soluble. Therefore, ~75% of Aβ variants maintained or increased the peptide’s propensity to aggregate in yeast cells.

To verify that our deep mutational scan accurately measured variant effects on aggregation, we tested six Aβ variants, G38F, K16V, A42V, F19Y, L17S and L34R, that spanned the solubility score range in a low-throughput validation assay. The growth rate of methotrexate-treated yeast expressing each Aβ variant was measured and compared to the aggregation propensity scores (**Figure 2C, S1B**). We found that the low-throughput assay results strongly correlated with the solubility scores derived from deep mutational scanning (R^2^ = 0.98). Thus, our assay reliably measured Aβ variant aggregation propensity.

To explore the effects of each amino acid substitution on Aβ aggregation, we created an Aβ sequence-aggregation map (**Figure 2D**). Substitutions to aspartic acid and proline were most associated with Aβ solubility, as evinced by their median scores of 0.64 and 0.56, respectively (**Figure S2A**). Conversely, the most aggregation-associated substitutions were hydrophobic tryptophan and phenylalanine, with scores of −0.60 and −0.51, respectively. Moreover, hierarchical clustering of all 791 solubility scores by amino acid revealed that hydrophobic substitutions, except alanine, clustered together and were associated with greater aggregation than other classes of substitutions.

Next, we characterized each position in Aβ based on its mutational profile. Hierarchical clustering of variant solubility scores by position identified six distinct clusters (**Figures 2E-F; S2B-C**). In cluster 1, comprising positions 17-20, 31-32, 34-36, 39 and 41, substitutions tended to decrease Aβ aggregation compared to substitutions in other clusters (cluster 1 mean solubility scores = 0.64, all other clusters = −0.28; **Figure S2D**). In cluster 1, even substitutions to hydrophobic amino acids slightly decreased aggregation (mean solubility score = 0.17). The effects of substitutions in cluster 2 were similar to but less extreme than in cluster 1. Both clusters 1 and 2 are largely comprised of hydrophobic positions in the wild type Aβ sequence. Indeed, 80% of Aβ positions with hydrophobic wild type residues are in clusters 1 and 2. In stark contrast, within clusters 4, 5 and 6, hydrophobic substitutions generally increase protein aggregation (all mean: −0.15, −0.12 and −0.45; hydrophobic means: −0.29, −0.65, and − 1.04). Cluster 3 contains only two positions, 37 and 38. Here, every substitution except proline increased aggregation (all mean: −0.99, hydrophobic mean: −1.56). Given that cluster 1 is characterized by hydrophobic positions where hydrophilic substitutions profoundly decreased aggregation, we suggest that this cluster defined buried β-strands in the Aβ sequence.

Next, we compared our solubility scores to previous alanine and proline scans which reported Aβ_40_ fibril thermodynamic stability *in vitro* (ΔΔG). ΔΔG values were determined by measuring variant Aβ monomer concentration remaining in solution after fibril formation reached equilibrium (Williams *et al.* 2004; Williams, Shivaprasad, and Wetzel 2006a). We found that the effects of proline substitution in our assay were correlated with proline ΔΔG values (R^2^ = 0.40), while the effects of alanine substitutions in our assay were less correlated with alanine ΔΔG values (R^2^ = 0.17; **Figure 3A**). In our alanine and proline comparisons, we found the greatest correlation at positions 17-20 and 31-32, where substitutions decreased aggregation in all studies (**Figure S3**). At positions 1-9 alanine or proline substitutions had smaller effects on aggregation. The most notable disagreement between studies was for alanine substitutions at positions 37 and 38. In our assay, alanine substitutions caused a profound increase in aggregation, whereas the *in vitro* alanine scan showed the opposite effect.

**Figure 3.**
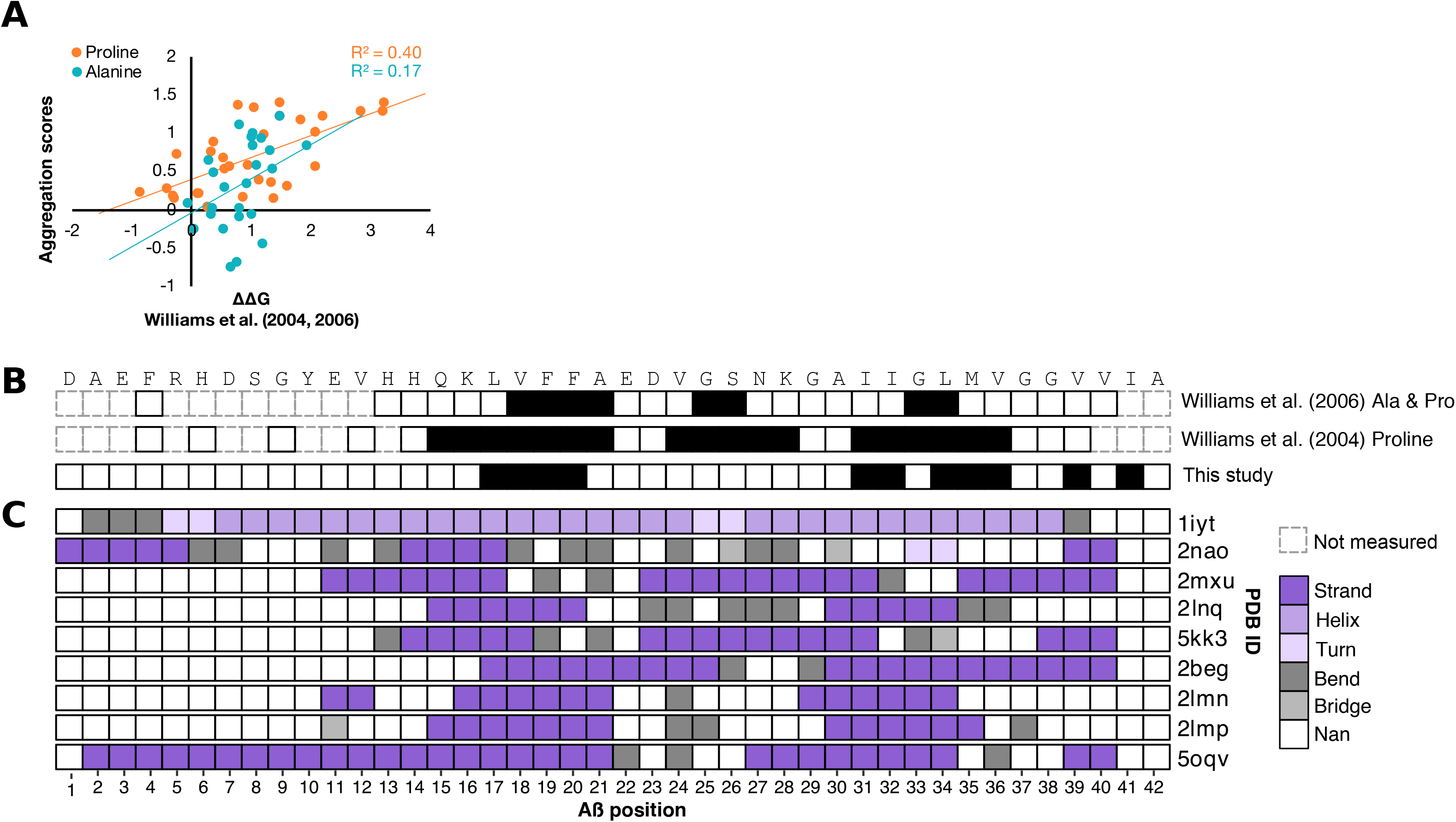
Comparison of yeast cell-based solubility scores to *in vitro* aggregation measurements and Aβ structural models. The scatterplot shows the correlation between our solubility scores (y-axis) and two single amino acid scans that measured the effect of proline or alanine variants on the thermodynamic stability of aggregates, relative to wild type (ΔΔG) (**A; Figure S3**). The first two tracks show unmeasured (gray) and the Aβ buried β-strand positions (black) suggested by proline scanning alone, or by proline and alanine scanning together (Williams *et al.* 2004; Williams, Shivaprasad, and Wetzel 2006a). The third track shows positions with the greatest increase in solubility when mutated in our large-scale mutagenesis study (**B**). The next nine tracks show the secondary structure of nine models of Aβ aggregate structure for each Aβ position (x-axis; **C**). The Aβ wild-type sequence is shown at the top.

We also compared the buried β-stand positions we proposed based on our deep mutational scanning data to β-stands proposed based on the alanine and proline scans, finding some concordance (**Figure 3B**). The single amino acid scans identify three regions that disrupt fibril elongation thermodynamics when mutated. The regions include positions 15-21, 24-28, and 31-36 for the proline scan and positions 18–21, 25–26, and 32–33 for the combined alanine and proline scans (Williams *et al.* 2004; Williams, Shivaprasad, and Wetzel 2006b). Given the generally highly disruptive nature of proline substitutions (Gray *et al.* 2017), it is not surprising that the proline scan would nominate many positions. Our deep mutational scan, on the other hand, does not reveal a central β-strand or strong decrease in aggregation with alanine or proline substitution from positions 24-28. We speculate that this difference is due either to the distinct experimental approaches used or to the different Aβ species (Aβ_40_ vs. Aβ_42_).

## Discussion

We used deep mutational scanning to characterize 791 Aβ variants in a yeast-based aggregation assay. Proline and aspartic acid substitutions were most disruptive of Aβ aggregation, while tryptophan and phenylalanine increased aggregation most. Additionally, we used unsupervised clustering to determine the regions of Aβ most important for aggregation. We conclude that these regions are most likely to form buried β-stands, which are necessary for aggregation and sensitive to amino acid substitutions (Jahn *et al.* 2010; Abrusán and Marsh 2016). These include positions 17-20, 31-32, 34-35, 39 and 41. While other positions could also form β-stands, the positions in cluster 1 are most likely to form the buried cores of Aβ aggregates in our cell-based assay.

Due to the noncrystalline nature of Aβ fibrils, traditional techniques such as X-ray crystallography and solution-state NMR cannot be used to solve Aβ’s aggregate structure. Instead, structural models have been developed by amassing constraints, such as the direction and register of β-sheets. For example, solid-state nuclear magnetic resonance studies suggest that Aβ fibrils are parallel, in register β-sheets (Benzinger *et al.* 1998; Gregory *et al.* 1998; Antzutkin *et al.* 2002; Tycko 2011). Many of these structural models are problematic because they are generated from constraints derived from *in vitro* experimental data, which may not be representative of *in vivo* conditions.

Given that we collected large-scale mutagenesis data in a cell-based system, we examined how our results compared to structural models of Aβ fibrils. Some models such as 1IYT (Crescenzi *et al.* 2002) and 2NAO (Wälti *et al.* 2016), showed very little to no overlap with either our proposed buried β-strands or those proposed by Williams *et al.* (2004, 2006) (**Figure 3C**). Other models contained three β-strand regions reminiscent of those suggested by Williams *et al.* (2004, 2006): 2MXU (Xiao *et al.* 2015), 5KK3 (Colvin *et al.* 2016), and 5OQV (Gremer *et al.* 2017). Yet other models propose β-strand patterns more similar to ours. These include 2BEG (Lührs *et al.* 2005), 2LNQ (Gremer *et al.* 2017), 2LMP and 2LMN (Lu *et al.* 2013). Since our β-strands were derived from data gathered in a cell-based assay, we hypothesized that they would be most consistent with structural models based on *in vivo*-derived fibrils. Indeed, the 2LMP and 2LMN models were based on fibrils seeded from plaques isolated from the brains of individuals afflicted by Alzheimer’s disease. Moreover, every model besides 2LMP and 2LMN was constructed using NMR or cryo-EM data from laboratory grown fibrils. That these models are less concordant with our cell-based mutational data suggests that there are important structural differences between *in vitro* and *in vivo* derived fibrils.

Deep mutational scanning data could contribute to the investigation of Aβ fibril structure beyond the analysis of existing models we present. For example, others have used site-saturation mutagenesis and deep mutational scanning data to evaluate proposed structural models (Bajaj *et al.* 2008; Khare *et al.* 2019). Additionally, deep mutational scanning data have now been used to generate distance constraints for the prediction of tertiary protein structure (Schmiedel and Lehner 2018; Rollins *et al.* 2018).

In summary, we used deep mutational scanning to elucidate the effects of amino acid substitutions on Aβ aggregation in a cell-based model. We used these large-scale mutagenesis data to propose positions critical for Aβ aggregation. Our results conflict with some previous *in vitro* reports of the effects of substitutions on Aβ aggregation and with some models of Aβ fibril structure. This outcome highlights the difficulties of studying protein aggregation and emphasizes the potential utility of *in vivo* or cell-based models. We suggest that deep mutational scanning of other aggregation-prone proteins such as α-synuclein or transthyretin could help reveal the relationship between sequence, structure and aggregation.

## Supporting information

SuppTable

SuppText

## Acknowledgements

This work was supported by the National Institute of General Medical Sciences (grant R01GM109110 to D.M.F.) and the Alzheimer’s Association. D.M.F. is a CIFAR Azrieli Global Scholar. V.E.G. was supported by a National Science Foundation Graduate Research Fellowship and a NHGRI Ruth L. Kirschstein National Research Service Award (T32HG000035). N.H. was supported by a NCI Ruth L. Kirschstein National Research Service Award (F30CA236335-01).

## Author Contributions

D.M.F. conceived of the project. D.M.F., V.E.G. and K.S. designed experiments. N.H designed the Aβ library. V.E.G., K.S., F.N.K., J.S. executed experiments. V.E.G., M.W. and F.N.K. analyzed data. V.E.G. and D.M.F. wrote the manuscript.

---

**Figure S1A-B.**
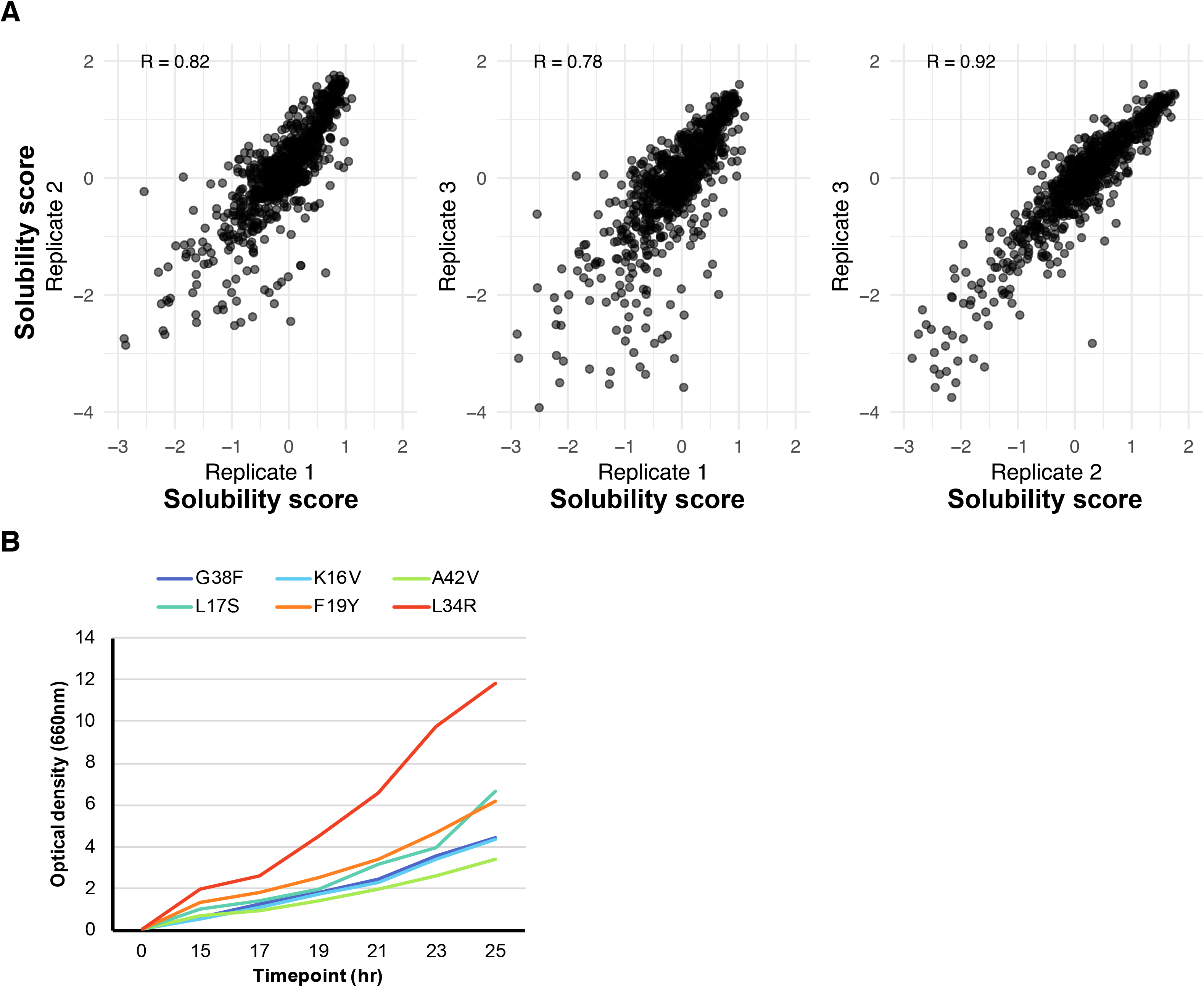
Validation of Aβ deep mutational scanning data. Scatter plots show correlations between variant solubility scores from three biological replicates. Pearson’s correlation coefficients (R) range from 0.78 to 0.92 (**A**). Six Aβ variants (G38F, K16V, A42V, F19Y, L17S and L34R) were tested in a low-throughput yeast-based aggregation assay, separately. The growth curves show spectrophotometer optical density (OD; y-axis) of yeast expressing a single variant over a 25 hour period (x-axis; **B**).

**Figure S2A-D.**
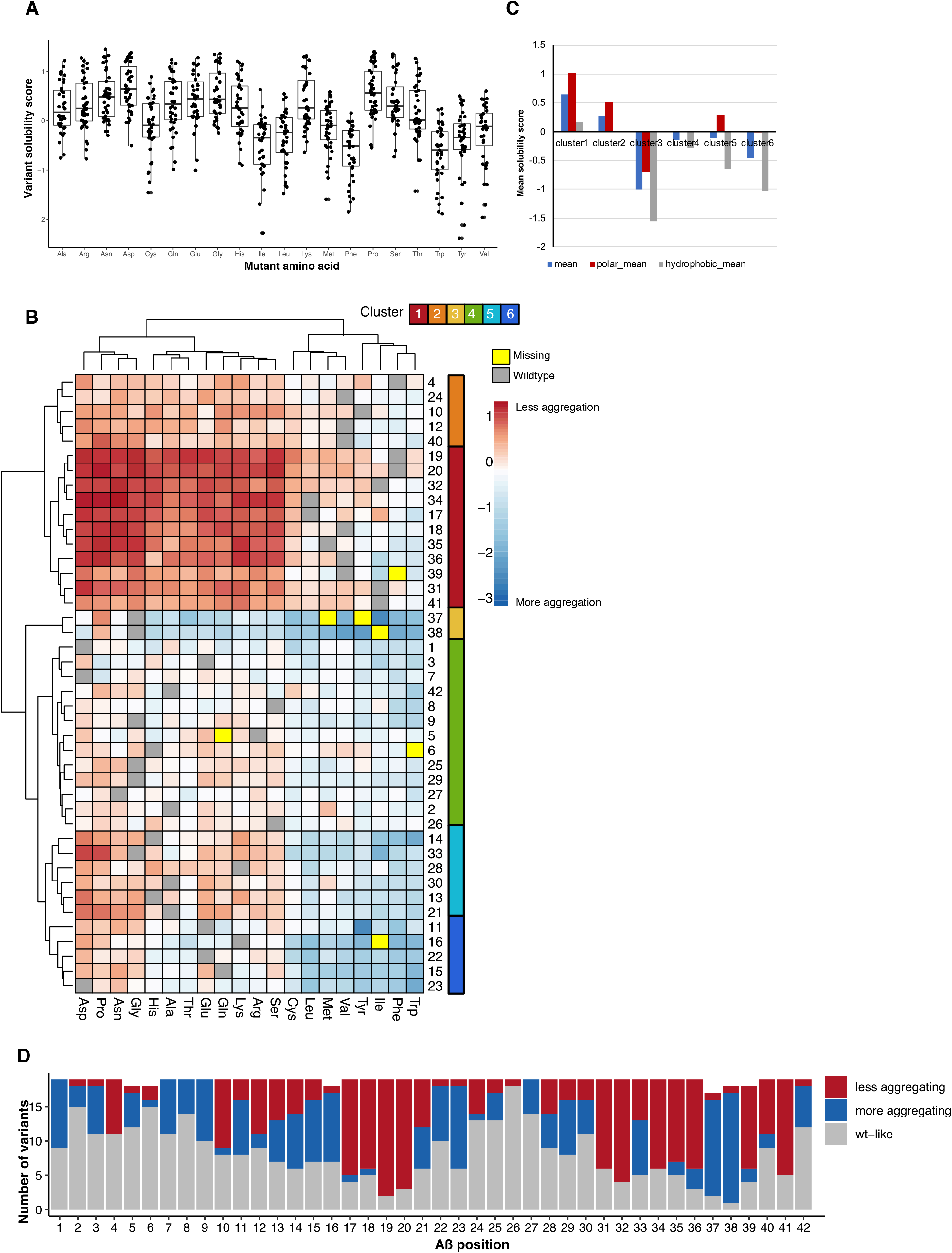
Large-scale mutagenesis summary and hierarchical clustering. The boxplots show the distributions of solubility scores (x-axis) observed for each substitution (y-axis; **A**). For example, the leftmost boxplot shows the solubility score for every alanine variant in our large-scale mutagenesis experiment. The heatmap contains the same information as **Figure 2D**, however hierarchical clustering has been applied to Aβ positions (y-axis; **B**). The six resulting clusters are shown in the annotation track to the right of the heatmap. The bar graph shows the mean solubility scores for each cluster identified by clustering in blue. In addition, mean solubility scores for polar and hydrophobic variants are in red and light gray, respectively (**C**). The stacked bar graph shows the number of variants (y-axis) that are wild-type-like, more and less aggregation-prone than wild-type at each Aβ position (x-axis; **D**).

**Figure S3.**
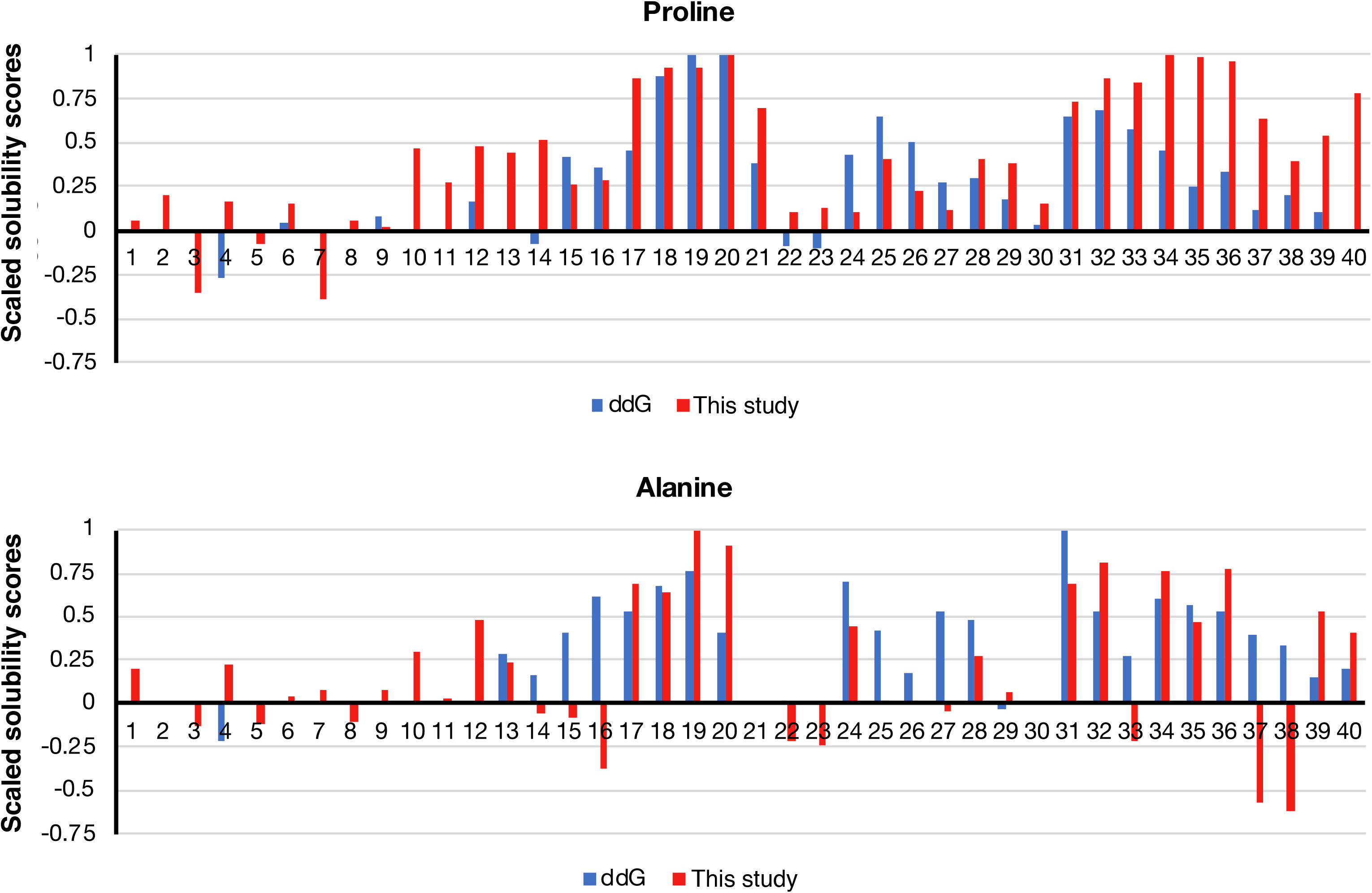
Position-wise comparison of our cell-based solubility scores with *in vitro* ΔΔG scores from(Williams *et al.* 2004; Williams, Shivaprasad, and Wetzel 2006a). Solubility scores and ΔΔG scores were scaled from 0 to 1, by dividing each score by their maximum values. The top panel shows the scaled solubility (red) and ΔΔG scores (blue; y-axis) for proline substitutions at each Aβ position (x-axis). The bottom panel shows alanine substitution effects (x-axis).

